# Fine-scale movements and habitat use of fish in intermittent rivers: Behavioural insights from drying refuge pools

**DOI:** 10.64898/2026.04.07.716892

**Authors:** Jolle W. Jolles, Jakob Gismann, Anna Cornet Sanz, Núria Bonada

## Abstract

Intermittent rivers and ephemeral streams (IRES) are increasingly recognised as ecologically important freshwater systems, yet little is known about how fish behave during the critical disconnected-pool phase, when drying confines them to isolated refuge pools. We combined high-resolution orthomapping and depth reconstructions with repeated whole-pool observations and focal follows to quantify microhabitat use and fine-scale movements of fish in refuge pools of an intermittent Mediterranean river. Fish used only a small fraction of the available pool area and consistently preferred deeper, refuge-associated microhabitats. Body size strongly structured behaviour: fry and small juveniles concentrated in shallow margins and showed short, tortuous movements, whereas larger individuals occupied deeper, more structured areas, moved farther, and were more closely associated with refuges. These patterns were broadly similar in drying and non-drying pools and changed little as water levels declined. After rewetting, fish showed reduced activity and weaker depth-biased habitat use, revealing that drying history leaves carry-over imprints on behaviour even after water levels recover. Our results show that refuge pools are not homogeneous water bodies, but internally structured habitats whose fine-scale characteristics shape how fish cope with drying, underscoring their conservation importance in increasingly intermittent rivers.

## 1. Introduction

Most of the world’s rivers and streams periodically cease flow [1], making flow intermittency a globally widespread feature of freshwater systems. In Mediterranean regions, this pattern is particularly pronounced, with intermittent rivers being the dominant freshwater type [2,3]. Drying fundamentally alters river ecosystems by fragmenting channels into a mosaic of flowing reaches, disconnected pools, and dry riverbeds, with major consequences for habitat availability, loss of connectivity, and degradation of local abiotic conditions [4,5]. These intermittent rivers and ephemeral streams (IRES) contribute disproportionately to freshwater biodiversity due to their pronounced spatial and temporal heterogeneity [6–8]. Recognition of this ecological importance has driven a rapid expansion of IRES research over the past decade, establishing intermittency as a central theme in freshwater ecology [9–11]. As climate change and human-induced alterations of land and water use are expected to increase the extent, duration, and severity of intermittency worldwide [3,11–14], there is a growing need to understand how organisms persist under increasingly dynamic and fragmented aquatic conditions.

As key components of river food webs, fish play a central role in structuring aquatic communities, and their loss can have cascading effects on ecosystem functioning [6]. Because most fish cannot survive complete desiccation, persistence during drying critically depends on access to remaining aquatic refuges [15–18]. However, most of our understanding of fish ecology and behaviour comes from perennial systems, with comparatively little attention given to the role of intermittency [18]. In IRES, research on fish has focused primarily on documenting the consequences of drying for species composition, abundance, and persistence, including pronounced shifts in assemblages between permanent and intermittent reaches [19,20] and high mortality during prolonged dry periods [15,21]. Together, these studies provide important insights into the demographic impacts of habitat contraction [4,5,16,17]. Beyond assemblage-level responses, research has also examined how fish respond behaviourally to drying, particularly in terms of habitat use and movement. Studies of habitat use show that fish occupy space non-randomly, with consistent preferences for deeper habitats, especially under low-flow conditions, in both perennial and intermittent systems [3,22–24]. Movement studies have further shown that fish often respond to drying by relocating into refuges as flow declines and dispersing again when connectivity is restored [25–30]. Together, this work has advanced understanding of how fish respond to drying at broader spatial scales, yet behaviour during the critical disconnected-pool phase remains poorly understood.

Once surface flow ceases, fish in intermittent rivers become confined to disconnected pools that constitute their only remaining aquatic habitat. During this phase, dispersal opportunities are strongly reduced and populations may experience severe bottlenecks. Although the importance of this disconnected-pool phase for ecosystem processes and persistence in IRES is widely recognised [5,31], fish-focused research has remained limited and has mostly concentrated on differences among pools, such as variation in species assemblages, survival, and abiotic conditions linked to pool morphology or physiochemistry [32–34]. By contrast, much less is known about how fish use space and move within refuge pools once flow has ceased. Under these conditions, the entire pool effectively functions as a single mesohabitat, placing strong constraints on behavioural decisions. Fine-scale behaviours, such as depth use, spatial positioning, and movement dynamics, are therefore likely to have direct consequences for exposure to stressors, individual condition, and persistence. Empirical data capturing these within-pool behavioural processes remain scarce (but see e.g. [35]), limiting our understanding of how fish cope with drying during this critical phase. Moreover, even less is known about how behaviour within these pools changes as drying progresses or whether the experience of drying leaves lasting imprints on fish behaviour once water returns, questions that are critical for understanding not just survival during the dry phase but recovery and persistence across the full drying cycle

Here, we address this gap by combining high-resolution reconstructions of pool morphology with repeated whole-pool observations and focal follows to quantify how fish use space and move within refuge pools during the disconnected-pool phase in an intermittent river. This approach allows us to characterise behaviour across multiple scales - from overall pool-level space use to within-pool habitat associations and fine-scale individual movement dynamics - while also examining how these patterns differ among fish of different sizes and life stages, given that habitat use and movement are often strongly size- and age-dependent [22,36–39]. We further examine three ecologically relevant contrasts: differences between fish in seasonally drying and nearby non-drying pools, changes in behaviour as water levels progressively decline, and behavioural carry-over effects before versus after drying under comparable water levels. Our study provides a close-up, in-situ view of fish behaviour within refuge pools that is rarely considered in freshwater research and offers insights into how animals cope with habitat loss and environmental change.

## 2. Material and Methods

### (a) Study system

Mediterranean intermittent river are among the most hydrologically stressed freshwater systems worldwide [2,3,40,41]. Our study focuses on the Borró, an intermittent stream in the foothills of the Pyrenees in north-eastern Catalonia, Spain (figure 1). Like many Mediterranean streams, it shows strong seasonal variation in water availability, with continuous flow during short wet periods but extended dry phases in summer when surface flow ceases and only disconnected pools remain [2]. We selected the Borró as a model system after extensive surveys in north-eastern Catalunya, as it exhibits pronounced drying dynamics, persistent fish populations, clearly defined refuge pools, and minimal human disturbance. The river forms part of the Alta Garrotxa network, a protected area of high ecological quality within the European Natura 2000 system [42]. We focused on a 2.3 km study reach in the upper part of the river’s seasonally wetted section, which we have monitored in detail for changes in water availability and fish presence over the past four years. This reach includes a longer intermittent section that dries almost completely - retaining less than about 10% of its wetted area in the driest summers - and a smaller upstream bedrock section that remains perennial year-round (figure 1). Further upstream, the river typically dries completely over several kilometres during summer.

**Figure 1.**
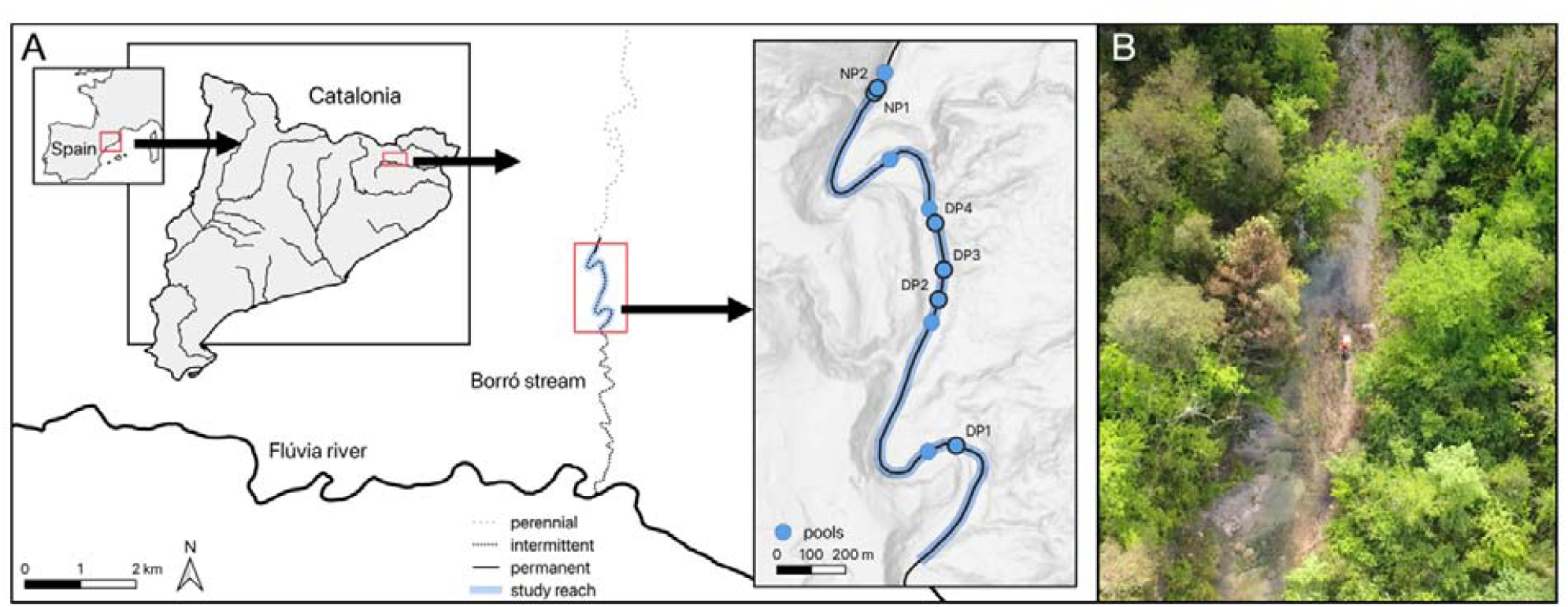
(A) Location of the study reach in the Borró stream (Fluvià catchment) in north-eastern Catalonia, Spain. The inset panel shows disconnected pools observed at the height of summer the preceding year, with focal pools labelled as DP (drying pools) or NP (non-drying pools). (B) Aerial view of the lower end of the study reach (near DP1) prior to the start of data collection.

### (b) Study species

Our study focuses on the Mediterranean Barbel (*Barbus meridionalis*), a small-bodied cyprinid endemic to north-eastern Spain and southern France [43]. The species is resident year-round and thus experiences the full range of drying and rewetting cycles, showing notable tolerance to intermittency [43,44]. Within the study reach, *B. meridionalis* is the dominant native fish and co-occurs mainly with the Catalan chub (*Squalius laietanus*) [42]. Although still relatively widespread, populations of *B. meridionalis* have declined due to flow alteration, habitat degradation, and introduced fishes [43,45,46], and the species is currently listed as Near Threatened on the IUCN Red List [47].

### (c) Study pools

For the present study, we selected six pools within the study reach (figure 1; table S1). Four pools were located in the intermittent section and showed strong seasonal declines in depth and size but remained wet through the driest summers, whereas two pools were located in the perennial section and maintained a stable depth year-round, hereafter referred to as drying and non-drying pools respectively. Pools that could not be fully observed from a single vantage point were divided into multiple observation zones of roughly equal size. In two elongated pools (DP2 and DP4), dense vegetation and steep banks prevented full visual coverage, so a single representative mid-section was selected for observation. For each zone, a fixed observation point was selected that provided an unobstructed view of the pool. All pools had clear water overall, with maximum depths ranging from 54-167 cm. Although absolute fish abundance could not be quantified, all pools had at least 50 fish and representation of all age classes and broadly comparable densities.

### (d) Pool mapping and depth survey

We created accurate two-dimensional maps of each pool with fine-scale depth information before the start of monitoring. First, we captured overlapping aerial images from a fixed height using a drone (DJI Mavic Mini), which were merged into a geo-referenced orthophoto using WebODM. This produced a composite image for each pool with a spatial accuracy of approximately 1 cm (figure 2A). Second, we used each orthophoto to produce simplified schematics showing the water perimeter and distinct features such as major rocks, fallen trees, or substrate changes (figure 2B), which aided us orient ourselves during observations. When underwater features were not visible due to surface reflection, their locations were mapped in situ using a laser rangefinder (AOFAR HX-700N) or measurement tape. Each schematic was overlaid with a 1 × 1 m grid and used as a reference for subsequent observations (see below). Third, we collected detailed depth and refuge information by wading through each pool, taking depth measurements roughly every 50 cm and at major depth transitions to 1 cm precision (figure 2B), and marking the location of underwater refuges such as large stones and overhanging banks. Depth points and refuge polygons were recorded live on a laptop using a custom Python tool that allowed interactive annotation on the pool schematic. One depth point per pool served as a reference to track water-level changes over time.

**Figure 2.**
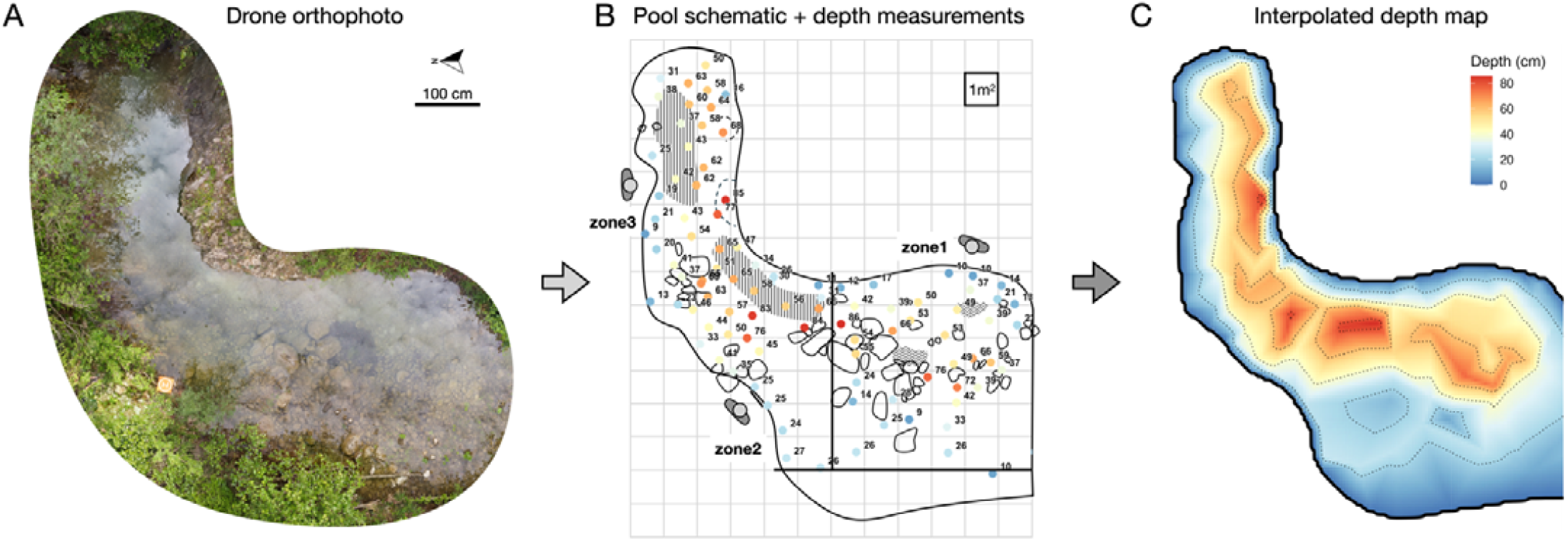
Workflow used to reconstruct pool bathymetry. (A) Orthophoto of a refuge pool (DP3) generated from drone imagery. (B) Simplified pool schematic used for pool observations, showing the 1 m^2^ grid and depth measurements (cm) collected throughout the pool with point colours corresponding to depth values. (C) Interpolated depth map used for our analyses.

### (e)Fish monitoring and behavioural observations

#### (i) Overview

Fish monitoring consisted of repeated, high-resolution observations of fish’s spatial distribution and behaviour across the drying-rewetting cycle. We combined two complementary methods: (1) whole-pool grid observations, where we recorded fish densities across predefined grid cells, and (2) focal follows, where we tracked pseudo-randomly selected individuals for one-minute periods throughout the pools. Monitoring took place over two months, starting at the end of May, allowing us to capture temporal changes in pool hydrology. We observed one to four pools per field day, scheduled so that every pool was monitored approximately once per week. Observation sessions were pseudo-randomly distributed throughout the day between 09.00 and 17.30 h to capture diel variation (figure S1). In total we conducted 53 pool sessions across 20 field trips, yielding 106 full grid observations and 374 focal follows. Fish were assigned to five size classes: s1 (fry; <1 cm), s2 (1-3 cm), s3 (3-6 cm), s4 (7-12 cm), and s5 (>12 cm). Most sessions occurred under sunny (69.8%) or partly sunny (15.1%) conditions, with fewer conducted under cloudy (7.5%) or overcast (7.5%) skies.

#### (ii) General protocol

Each observation session began with the observer slowly approaching the fixed observation platform, setting up equipment, and remaining still for five minutes to minimize fish disturbance. Monitoring sessions consisted of one grid observation, followed by 5-10 focal follows, and concluded with a second full grid observation. For pools with multiple zones, these steps were completed sequentially in the same order for each zone. After each session, the observer recorded meta-information including weather, which grid cells were in direct sunlight, a depth measurement at the reference point, and any additional relevant notes.

Data were recorded either on printed pool schematics or digitally using an iPad and the Sketchbook app. Observers consistently wore polarized sunglasses to reduce surface glare and used short-range binoculars (Vortex Diamondback 8×42 fitted with polarizing lens covers) to improve detection of small or distant fish. When a surface film or algae layer was present, it was removed with a hand net and observations were postponed until later in the day. Consistency in assigning individuals to size classes was ensured through repeated training among observers. When uncertain during observations, fish were compared with nearby individuals to recalibrate size estimates.

#### (iii) Grid observations (GO)

Grid observations involved systematically scanning each zone and recording fish occurrence across all grid cells. For each grid cell, fish presence was recorded by drawing dots on the pool schematic. Counts were recorded categorically using one to three dots (representing 1-3, 4-10, and >10 individuals, respectively), and five distinct colours were used to represent the different size classes. Each full grid survey lasted approximately 5 s per wet cell, during which cells were scanned sequentially in pseudo-random order (starting in different corners and alternating between row, column, clockwise, or counter-clockwise sequences). When individual fish moved between adjacent cells during a scan or were recognized from a neighbouring cell, they were not recorded again to avoid double-counting. This procedure yielded a structured, high-resolution representation of fish presence and spatial distribution across each pool.

#### (iv) Focal follows (FF)

Focal follows involved continuously tracking individual fish for one minute while drawing their trajectories live on the pool schematic and recording their size class. Fish were pseudo-randomly selected to obtain approximately equal numbers across size classes and from different areas within each pool. Trajectory endpoints were marked with a small arrow indicating swimming direction. Shoaling behaviour was recorded by scoring the number of conspecifics within four body lengths of the focal fish [48] and categorised into four classes: 0, 1-2, 3-9, or ≥ 10 fish.

### (f) Data processing

#### (i) Depth and refuge data

Depth measurements and refuge polygons extracted from pool maps were first transformed into a common two-dimensional coordinate system aligned with the observation grid, such that grid positions corresponded to real-world distances expressed in centimetres. We then computed a depth map for each pool at 10 cm resolution (figure 2C) using a custom R function based on the interpolation algorithms of the akima package [49]. All maps were visually checked for consistency with the original depth-point data and known pool topography. At the start of the study, pools covered on average 86.5 m^2^ of wetted area (range: 49.6-133.9 m^2^), and 22.2 % of grid cells contained refuge features (range: 8.9-38.3 %).

Using the reference depth measured during each monitoring session, we estimated the session-specific maximum water depth and calculated the change in water level relative to the reference state. This allowed us to generate adjusted depth maps with updated water-edge positions and to compute corresponding changes in surface area (figure S3). From each session-specific adjusted depth map, we then derived for every 1 m^2^ grid cell the presence of water, mean and maximum depth, and refuge presence.

Because pools differed in depth and drying pools became progressively shallower during summer, depth values were expressed relative to each pool’s maximum depth at the start of the study. Relative depth was calculated as the difference between the current maximum pool depth and the local depth, scaled by the deepest mean depth observed in that pool at the start of the study. Values close to 0 therefore indicate locations near the deepest part of the pool, whereas higher values indicate progressively shallower positions. This approach accounts for declining water levels while preserving the relative bathymetric position of locations, allowing direct comparison across pools and over time. For example, a cell that is 80 cm deep in a pool with a maximum depth of 100 cm has a relative depth of 0.2; if water level declines by 20 cm, the same cell becomes 60 cm deep while pool maximum depth becomes 80 cm, and its relative depth remains 0.2.

#### (ii) Observational data

Spatial data extraction from digital pool observations was performed with a custom Python workflow while data of the manual observations was scored by hand. First, coordinates of all grid cells were defined for each pool image. Second, for grid observations, we extracted the coordinates and colours of all recorded dots and converted them into categorical abundance values per size class for each grid cell. Third, focal-follow trajectories were reconstructed using the manual tracking function of ATracker [50] on the grid pool image. All point coordinates were then converted to grid-relative centimetre units and standardized to equal time intervals. Spatial coverage of grid observations and focal-follow trajectories within a representative pool is illustrated in figure S2.

For each focal follow, we recorded the social state (if a fish was shoaling during any part of the follow) and calculated the proportion of time spent shoaling, total distance moved, maximum displacement from the point of origin, and path straightness. Path straightness was quantified from the relationship between maximum displacement and total distance moved, with higher values indicating more directional trajectories and lower values more tortuous paths (see figure S3). We additionally calculated the mean and maximum depth along the trajectory and the number of grid cells crossed that contained a refuge. For grid observations, we calculated pool-level occupancy by dividing the number of cells occupied by each size class by the total number of wet cells. Because 93% of occupied cells fell within the two lowest abundance categories, grid observations were analysed as presence/absence rather than categorical abundance.

#### (iii) Environmental data

Because environmental conditions within pools vary across space and time and may influence fish activity and habitat use, we quantified sun exposure and water temperature as contextual variables. For each session, we digitised which wet grid cells received direct sunlight (hereafter sun exposure, figure S4A) and calculated the proportion of wet cells in the sun. Across the 53 unique sessions, 13 had no direct sun exposure, whereas the remaining sessions spanned the full range from low to complete exposure (figure S4B). Water temperature was recorded at 1 min intervals throughout the summer using a HOBO logger placed at approximately 40 cm depth in one randomly selected pool of each type (DP1 and NP1), attached beneath a large stone to avoid direct solar heating. These temperature series were used as representative values for the other pools of the same type.

To place pool-level environmental conditions in a wider temporal context and characterise seasonal changes associated with drying and rewetting, weather data were obtained from the Meteorological Service of Catalonia (Meteocat), including daily precipitation and average air temperature. Because the field site lies approximately midway between the two nearest stations (Banyoles and Olot), values from both stations were averaged. During the study period, mean daily air temperature ranged from 18.5 to 29.2 °C and water temperature from 14.5 to 25.9 °C, while total rainfall was 68.5 mm, concentrated across six rainy days (>1 mm, figure S5).

### (g) Data analysis

We used linear mixed-effects models to quantify how habitat structure, environmental conditions, body size, and seasonal changes in water availability shaped fish distribution and behaviour within refuge pools. Analyses were conducted at four levels: pool-level space use, within-pool habitat associations, individual movement and social behaviour, and temporal changes during seasonal drying and rewetting.

Pool use was quantified as the proportion of wet grid cells occupied during each observation session and analysed using linear mixed-effects models with pool type as a fixed effect and pool identity and date as random effects. Diel variation was tested by comparing models including linear versus quadratic effects of time of day. To assess environmental drivers, sun exposure (proportion of wet grid cells in direct sunlight) and water temperature were included as fixed effects in a separate model with the same random-effects structure.

Within-pool habitat use was analysed at the grid-cell level using binomial generalized linear mixed-effects models with cell occupancy (presence/absence) as the response. A size-collapsed model included relative depth, refuge presence, sun exposure, pool type, and the depth × pool type interaction as fixed effects, with random intercepts for pool and observation replicate. To assess size-dependent patterns, a size-explicit model included size class and its interactions with habitat variables. Differences in explanatory power were evaluated using marginal R^2^. Patterns from grid-cell observations were corroborated using focal-follow data: mean and maximum relative depth were analysed with linear mixed-effects models, and the probability of entering refuge cells with a binomial mixed-effects model, each including size class and pool type as predictors.

Individual movement and social behaviour were analysed based on the 1-min focal follow data. Maximum displacement, movement speed, path straightness, and turning rate per unit distance were analysed with linear mixed-effects models, and shoaling tendency (proximity to conspecifics) with a binomial mixed-effects model. Models included size class and pool type as fixed effects and pool and sampling session as random effects. Movement speed was expressed in body lengths per second. Turning-rate analyses excluded focal follows with very low displacement (<25 cm), as estimates are unreliable for near-stationary trajectories; results were qualitatively unchanged when including all observations.

Seasonal effects were analysed in two steps. First, drying dynamics were quantified by modelling maximum pool depth as a function of relative day using linear mixed-effects models with pool as a random effect. Behavioural responses during drying were assessed by refitting pool-level, habitat-use, and movement models using data from the drying phase, with relative day as the temporal predictor. For habitat use, interactions between relative depth and time tested for changes in depth preferences, and mean absolute and relative depth use were analysed separately. Second, to compare behaviour at similar water levels before and after drying, data were subset to early (≤ 7 June) and late (≥ 12 July) periods with comparable pool depths. Models were refitted with period as predictor, including interactions with pool type and, for movement, with size class.

All analyses were conducted in R 4.4.1 [53]. Mixed models were fitted using lme4 [54] and glmmTMB [55], with Gaussian or binomial error distributions as appropriate, and significance of fixed effects was assessed using likelihood-ratio tests. Model assumptions were evaluated using DHARMa [56]. When Gaussian assumptions were violated, models were refitted using a Gamma distribution with log link. Collinearity was assessed using variance inflation factors (VIF < 3). Model predictions were obtained using emmeans [57], and marginal and conditional R^2^ values followed Nakagawa and Schielzeth [58].

## 3. Results

### (a) Pool-level space use

Fish used a small fraction of the available wetted area of the pools per session (mean ± s.e. of wet grid cells: 18.3 ± 0.7 %, n = 106). This proportion was lower in drying than in non-drying pools (χ^2^ = 3.98, d.f. = 1, *p* = 0.046, figure 3A), even more so when considering only repeatedly used cells (5+ times) across all sessions (23.4% versus 41.8%), indicating that fish were consistently concentrated in a restricted subset of available habitat. Pool use also varied systematically over the day, with fish using most of the area around midday (quadratic model: χ^2^ = 14.14, d.f. = 1, *p* < 0.001), closely matching changes in sun exposure (χ^2^ = 12.06, d.f. = 1, *p* < 0.001, figure S6). In contrast, pool use did not change with daily fluctuations in water temperature (χ^2^ = 0.02, d.f. = 1, *p* = 0.891; figure S6).

**Figure 3.**
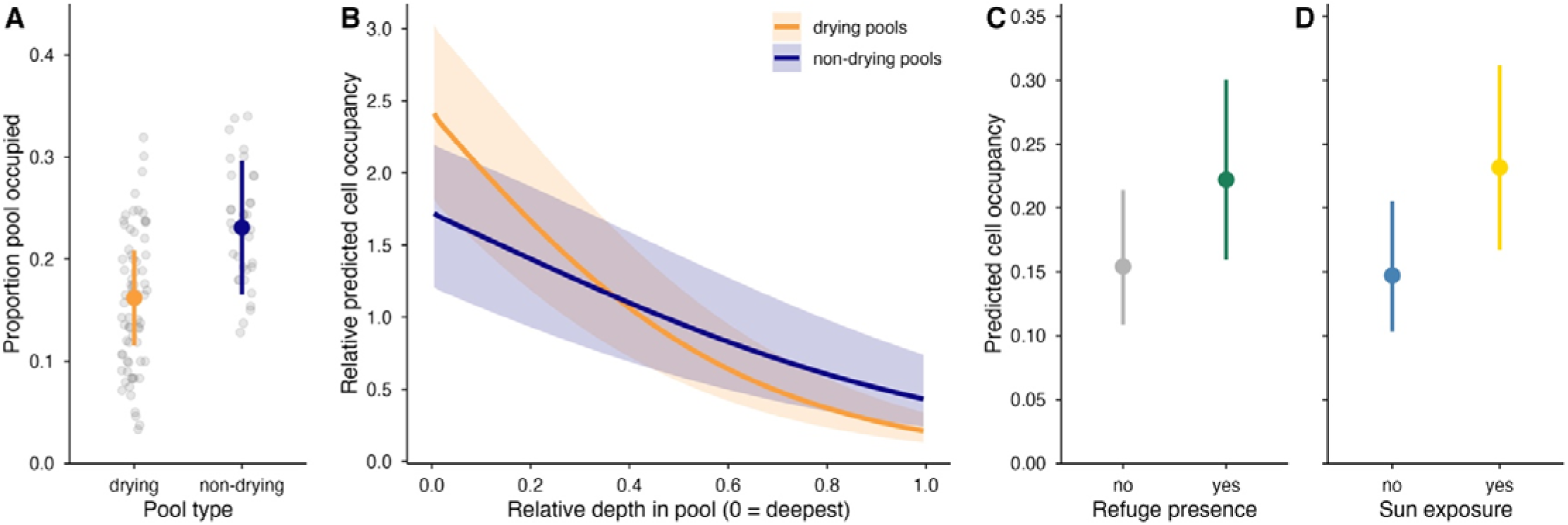
(A) Session-level pool occupancy in drying and non-drying pools. Points show individual observations and coloured points show model-predicted means. (B) Relative predicted cell occupancy as a function of relative depth in the pool (0 = deepest), shown separately for drying and non-drying pools. Predictions were scaled relative to an expectation of uniform use within each pool type. (C–D) Proportion of the pool occupied under different conditions, showing model-predicted probabilities that a grid cell is occupied when it contains a refuge or not (C) or is in the shade or in the sun (D). Error bars and shaded bands indicate 95% confidence intervals.

### (b) Within-pool habitat use

We next examined how space use was structured by local habitat features. Fish were considerably more likely to be found in deeper parts of the pools, and this depth-preference was stronger in drying than in non-drying pools (depth × pool type: χ^2^ = 22.24, d.f. = 1, *p* < 0.001, figure 3B). Refuge availability also had a strong effect on space use (χ^2^ = 53.42, d.f. = 1, *p* < 0.001), with fish more likely to occur in grid cells containing refuges than in open cells (figure 3C). Sun exposure further influenced space use, with fish more likely to occur in sun-exposed cells (χ^2^ = 72.60, d.f. = 1, *p* < 0.001; figure 3D).

Next, we examined habitat use separately for each size class rather than for fish occurrence overall. Including size class substantially improved model performance (marginal R^2^ = 0.29 versus 0.18) and revealed strong size-dependent differences in within-pool habitat use. Grid-cell observations showed that larger fish were much more likely to use deeper areas (χ^2^ = 234.0, d.f. = 4, *p* < 0.001, figure 4A) and, to a lesser extent, refuge cells (χ^2^ = 14.7, d.f. = 4, *p* = 0.005, figure 4B), whereas smaller fish were more likely to use sunlit cells (χ^2^ = 39.1, d.f. = 4, *p* < 0.001, figure 4C). Focal-follow data supported these patterns: larger fish occurred deeper on average (χ^2^ = 98.2, d.f. = 4, *p* < 0.001), with no difference between pool types (χ^2^ = 2.76, d.f. = 1, *p* = 0.598). They also used refuge areas more frequently (χ^2^ = 20.1, d.f. = 4, *p* < 0.001) and reached greater maximum depths, particularly in drying pools (χ^2^ = 11.8, d.f. = 1, *p* = 0.019).

**Figure 4.**
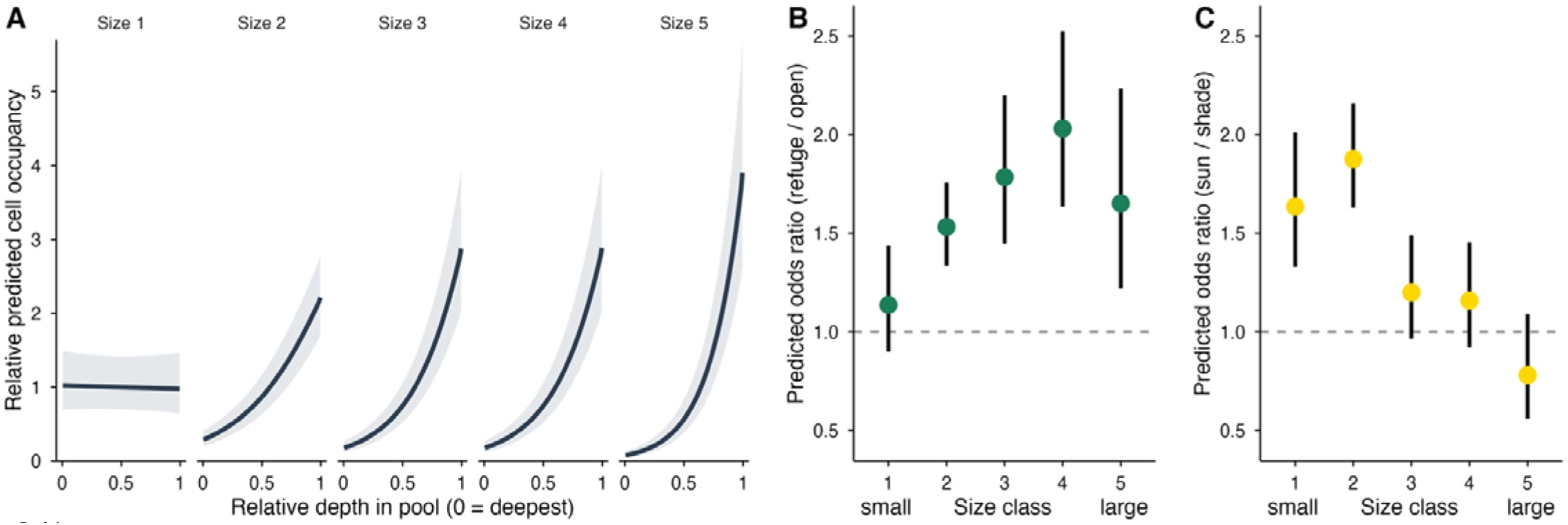
(A) Relative predicted cell occupancy as a function of relative depth in the pool (deepest = 0), shown separately for each size class. Predictions were scaled relative to an expectation of uniform use within each class. (B–C) Size-specific odds ratios for cell use in terms of (B) refuges versus open microhabitats and (C) sun-exposed versus shaded microhabitats. Points indicate model-predicted means and vertical lines show 95% confidence intervals, with predictions averaged over the remaining predictors.

### (c) Individual movement

We next examined individual movement patterns within the pools using focal-follow data. Movement differed strongly among size classes: larger fish moved further and showed greater maximum displacement (χ^2^ = 77.83 and 87.45, d.f. = 4, p < 0.001; figure 5A), whereas smaller fish were more active relative to their body size, moving faster (χ^2^ = 73.30, d.f. = 4, p < 0.001; figure 5B) and making more frequent directional changes (χ^2^ = 35.8, d.f. = 4, p < 0.001; figure 5C). Path straightness did not differ among size classes (χ^2^ = 3.93, d.f. = 4, p = 0.416). Fish moved in proximity to conspecifics on average 33.6% of the time, and this tendency varied among size classes (χ^2^ = 18.82, d.f. = 4, *p* < 0.001), with intermediate-sized fish being the most social (figure 5D). None of the movement metrics differed between drying and non-drying pools (all *p* > 0.15).

**Figure 5.**
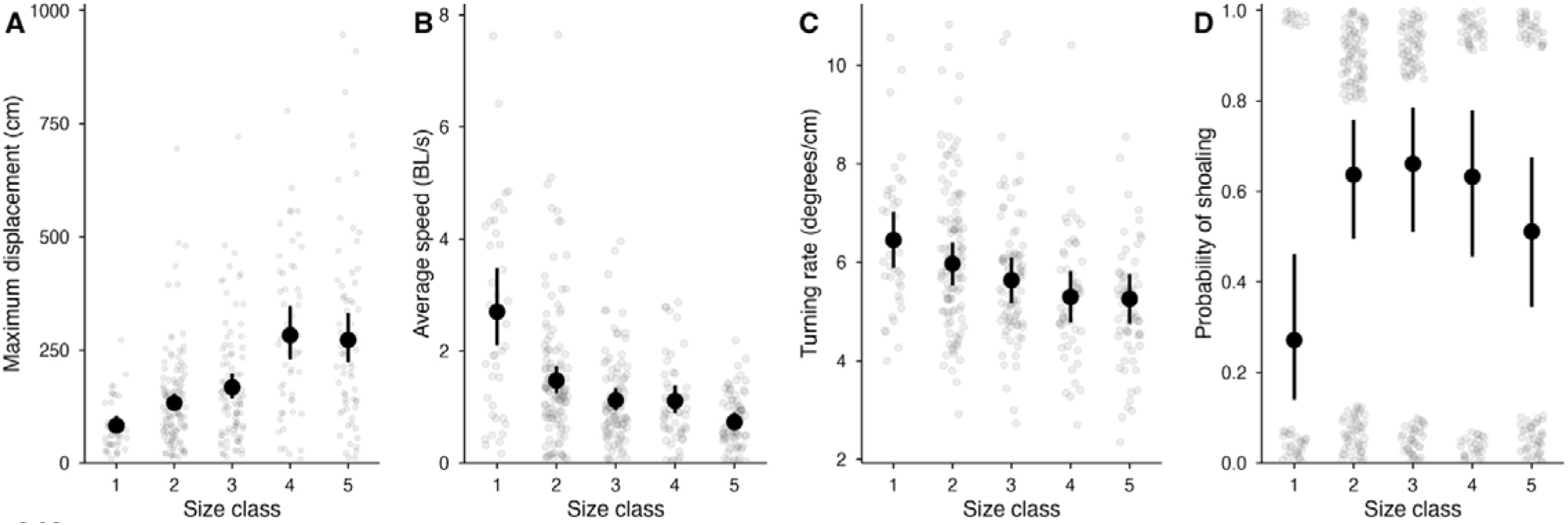
Size-dependent differences in fine-scale movement and social behaviour during 1-min focal follows. Grey points show individual observations and black points with error bars show estimated marginal means ± 95% CI. (A) Maximum displacement from the starting point (cm). (B) Average movement speed expressed in body length per second. (C) Turning rate per unit distance (focal follows with displacement >25 cm). (D) Probability of shoaling during the focal follow with raw binary observations (0/1) vertically jittered for visualisation. Size classes are ordered from smallest to largest.

### (d) Behavioural responses to seasonal drying

We next examined how fish behaviour changed over the seasonal drying period, both during progressive habitat contraction and following rewetting. During the first part of the observation period, drying pools contracted substantially (mean reduction 21.4%, range 6.7-44.4%) and became progressively shallower (-6 cm/week depth on average), whereas non-drying pools remained stable (figures 6A, S7–S8). Despite this marked reduction in habitat availability, fish showed little behavioural response. Pool use remained unchanged throughout the drying phase, both in absolute number of occupied cells (χ^2^ = 1.63, d.f. = 1, p = 0.201) and relative occupancy (χ^2^ = 1.31, d.f. = 1, p = 0.252). As pools became shallower, fish were found in progressively shallower water, but maintained similar positions relative to the deepest part of the pool (grid observations: χ^2^ = 0.09, d.f. = 1, p = 0.764; focal follows: χ^2^ = 0.21, d.f. = 1, p = 0.650; figure 6A). Movement behaviour also remained stable, with no change in total distance moved during focal follows (χ^2^ = 0.28, d.f. = 1, p = 0.600).

**Figure 6.**
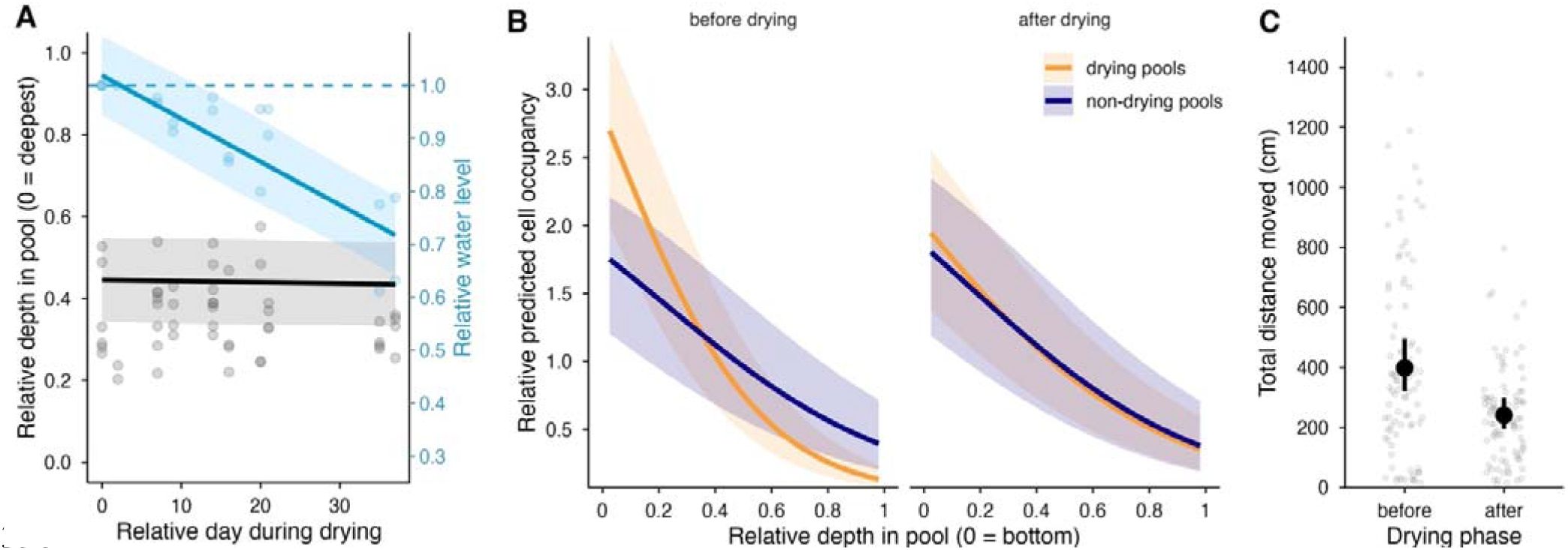
(A) Relative water level (blue, right axis) and mean relative depth of occupied cells (black, left axis) across the drying period. Points show mean observations per session. (B) Relative predicted within-pool habitat use across the depth gradient before drying and upon rewetting for drying (orange) and non-drying (blue) pools. Predictions were scaled relative to an expectation of uniform use within each pool type and period. (C) Total distance moved before and after the drying phase, with points showing individual observations. Shaded areas in A and B and error bars in C are 95% confidence intervals.

Following a short wetter period, pools refilled to depths comparable to those at the start of the season (figure S8), allowing comparison of behaviour before and after drying. Pool use did not differ between periods (χ^2^ = 0.18, d.f. = 1, p = 0.675). However, fish upon rewetting used relatively deeper parts of drying pools less than before the drying phase (relative depth × pool type × period: χ^2^ = 8.18, d.f. = 1, p = 0.004; figure 6B). Movement was also reduced after drying (χ^2^ = 8.95, d.f. = 1, p = 0.003; figure 6C), with no difference between pool types (χ^2^ = 0.30, d.f. = 1, p = 0.585) or among size classes (χ^2^ = 7.39, d.f. = 4, p = 0.117).

## 4. Discussion

Fish confined to refuge pools showed strongly structured fine-scaled patterns of habitat use and behaviour. Although responses to drying in intermittent rivers have been described at broader spatial scales [6,11,51], how fish use space and move within disconnected pools has remained largely unresolved. By combining high-resolution spatial reconstructions of pool bathymetry with repeated whole-pool observations and focal follows, we show that fish in refuge pools are far from randomly distributed: they consistently used deeper and refuge-associated microhabitats, with strong size-dependent differences in both habitat use and movement. These patterns were broadly similar across drying and non-drying pools and changed little during progressive habitat contraction, indicating that fish largely maintained their spatial organisation as pools shrank. By contrast, patterns following rewetting differed from those before drying, suggesting that drying history can leave a detectable imprint beyond immediate habitat conditions.

Both the pool grid-observations and focal follows showed that fish in refuge pools consistently preferred deeper areas and microhabitats containing refuges. Preferences for deeper habitats are well documented in freshwater fish, especially in permanent rivers where microhabitat selection has been studied extensively [22,24,36]. Far less work has addressed habitat use explicitly in intermittent rivers, but existing studies report broadly similar patterns at coarser spatial scales, with fish increasingly occupying pools and deeper habitats as flow declines [23,27,35,38]. Associations with refuges are likewise consistent with observations from both permanent and intermittent rivers [22,38,52,53]. Our findings extend this work by showing that these habitat preferences persist even after fish become confined to disconnected pools and remain stable across varying levels of drying. Sun exposure also influenced space use, with fish more likely to use sunlit cells and overall pool use peaking around midday. Given that water temperature had no effect on space use, this pattern more likely reflects improved prey detection visibility than thermoregulation. Thus, refuge pools should not be viewed as homogeneous water bodies once flow ceases, but as internally structured habitats in which fish organise their space use along fine-scale gradients.

Beyond the general preference for deeper areas within refuge pools, habitat use differed strongly among size classes, revealing pronounced ontogenetic differentiation during the disconnected-pool phase. The smallest size class, fry, was disproportionately associated with shallow pool margins, whereas progressively larger size classes increasingly occupied deeper, refuge-associated microhabitats. Given that the observed size classes ranged from fry of up to 1 cm to individuals exceeding 10 cm, this pattern points to marked behavioural structuring across life stages. Size- and age-related shifts in depth and cover use are well documented for stream fish in permanent rivers [22,24,39,53–55] and similar patterns have been reported in intermittent rivers at broader spatial scales [27,35,38]. Our results show that this structuring is maintained even under extreme spatial confinement, with size classes remaining strongly segregated within single refuge pools at fine spatial scales. Predation risk is often invoked to explain such size-dependent habitat use [22,37,38], but piscivorous predators were largely absent from our study pools and aerial predation was rarely observed. Predation may therefore contribute to the observed pattern, but is unlikely to be its sole driver. The stronger association of larger fish with deeper, more refuge-rich areas may instead reflect a preference for environmentally more stable microhabitats, as has previously been suggested at broader spatial scales [27,37]. Together, these findings show that refuge pools are not only internally structured, but also strongly size-segregated, with different life stages using distinct microhabitats during the most constrained phase of river intermittency.

Focal follows provided an additional, individual-level view of behaviour within refuge pools, revealing marked size-related differences in movement dynamics. At broader spatial scales, fish movements in intermittent rivers are typically described in terms of overall mobility within streams and shifts among habitats, including movements into pools during drying, dispersal toward perennial reaches, and dispersal following rewetting [26,28,30,56,57]. Our data show that, once confined to a single pool, movement behaviour remains highly structured. Larger fish moved further and sampled broader portions of the pool, whereas smaller fish showed more intensive localised movement, moving faster relative to their body size and changing direction more frequently. These patterns point to distinct movement strategies, with smaller, younger individuals engaged in fine-scale local exploration and larger, older individuals moving more broadly through the pool. At the reach scale, fish associated with pools and deeper habitats are often reported to move less than individuals occupying shallower habitats [22,52,58], typically interpreted as reduced need for displacement once suitable habitat is located. Our results show that this relationship differs at the within-pool scale: larger fish were both more strongly associated with deeper, refuge-rich microhabitats and more mobile within pools, suggesting that relationships between habitat use and movement are strongly scale-dependent. Such differences are likely to affect how individuals sample pool conditions and acquire spatial information as habitats contract, potentially influencing when and how they respond to local environmental change. By combining fine-scale behavioural observations with spatially explicit habitat data in situ, our study provides rare field-based insight into movement dynamics of early life stages and resolves within-pool variation that would remain hidden in broader-scale surveys.

Fish used only a small part of the available pool area, on average about one fifth of wetted cells, and despite substantial contraction of pool surface area this did not change as drying progressed. Likewise, fish did not retreat towards relatively deeper parts of pools as water levels declined, but continued to occupy similar relative positions and therefore occurred in shallower water overall. Movement behaviour also remained unchanged. Together, these results indicate that while progressive drying reduced the amount of habitat available, it did not alter how fish used space or moved within pools. Previous work has shown that fish relocate into pools as flow declines and remain there until connectivity is restored [23,26,27,51]. Our results extend this perspective by showing that, once confined, fish largely accommodate shrinking habitat without reorganising their fine-scale behaviour.

When pools refilled to comparable water levels following the drying period, fish moved less than before drying in both pool types, and only in pools that had previously dried did depth-biased habitat use weaken after rewetting, with depth preferences converging towards those observed in non-drying pools. The mechanism behind this carry-over effect remains unclear, but the finding itself is striking: drying history shaped how fish reorganised space use once water levels recovered, even when behaviour had changed little during drying itself. Notably, our study coincided with a mild summer and early rewetting, suggesting that carry-over effects may be stronger under more severe or prolonged drying. Whereas previous studies have documented effects of drying on fish assemblages and abundance [15,16,33], our results suggest that even mild drying can leave a detectable imprint on fine-scale behaviour, a distinction between acute and cumulative behavioural effects of environmental change that warrants further investigation.

Most previous studies of fish in intermittent rivers have inferred behavioural responses indirectly, relying on changes in presence, abundance, or assemblage composition [16,33], movements among pools or reaches inferred from mark-recapture or telemetry [26,27,51], or snapshot assessments based on snorkelling or electrofishing surveys [22,24,38]. Here, we go beyond these approaches by directly quantifying space use and movement within refuge pools during the disconnected-pool phase. By combining high-resolution orthomapping with repeated whole-pool observations and focal follows, we resolve behavioural variation at spatial and temporal scales that closely match the constraints fish experience during isolation. This provides a rare temporal perspective on fine-scale habitat use and movement during confinement, connecting within-pool behaviour to broader questions about how fish cope with habitat contraction.

Our study was necessarily limited to a single river and six focal pools. Although these pools were selected following long-term regional monitoring to capture representative conditions, broader sampling across rivers and pool types will be needed to better understand how variation in pool morphology, drying intensity, and environmental context shapes behaviour. A further limitation is that we did not quantify fish abundance or density within pools, and density-dependent effects could therefore in principle have contributed to some of the observed patterns, although fish numbers appeared broadly similar among focal pools and stable over time. Because our study coincided with a relatively mild summer and early rewetting, applying similar approaches across more extreme drying events will be particularly important for identifying when behavioural reorganisation becomes pronounced, whether thresholds emerge, and under what conditions refuge pools may cease to function as effective refuges and turn into ecological traps [17]. At the same time, confinement of fish within pools provided a valuable opportunity to follow behaviour repeatedly through time and relate it to progressive drying and rewetting. Future work should extend this perspective across stronger drying gradients and shallow-water environments, while linking behaviour more directly to coping mechanisms and survival under confinement.

Our study shows that fish confined to refuge pools do not use them as homogeneous water bodies, but as internally structured habitats in which space use and movement are finely organised along depth and refuge gradients and strongly differentiated by size. Behaviour changed little during progressive habitat contraction, yet drying history left a detectable imprint after rewetting, highlighting a distinction between acute and carry-over behavioural effects that has received little attention in intermittent river research. These findings also demonstrate what fine-scale, spatially explicit approaches can reveal that broader surveys cannot, contributing to our understanding of how animals adjust behaviour to persist in extreme and rapidly changing environments [59]. Ultimately, our results underscore that refuge pool internal structure and behavioural functionality matter for fish persistence during drying, with direct implications for how we assess and conserve refugia in increasingly intermittent river systems [60].

## Authors’ contributions

The study was conceived by J.W.J. Methodological development, including the design of grid observations and focal follows, was led by J.W.J. and J.G., with considerably further input from A.C.S. and helpful feedback from N.B. Drone surveys and orthomapping were conducted by J.W.J., depth measurements by J.W.J. and J.G., and pool schematics drawn by A.C.S. Behavioural field data were collected by A.C.S., J.G., and J.W.J., with A.C.S. contributing the majority of behavioural observations. J.W.J. processed the data, generated depth maps, computed the behavioural metrics, and performed all statistical analyses. J.W.J. wrote the manuscript. N.B. contributed to conceptual framing within the context of intermittent river ecology. All authors approved the final manuscript.

## Acknowledgements

We thank Emili García-Berthou and Frederic Bartumeus for early support that helped J.W.J establish the foundations of this research line. We are grateful to Galdric Mossoll for his dedication during the initial exploration and set-up of the field sites and data collection of their drying dynamics.

## References

1. Messager ML et al. 2021 Global prevalence of non-perennial rivers and streams. Nature 594, 391–397. (doi:10.1038/s41586-021-03565-5)

2. Bonada N, Resh VH. 2013 Mediterranean-climate streams and rivers: Geographically separated but ecologically comparable freshwater systems. Hydrobiologia 719, 1–29. (doi:10.1007/s10750-013-1634-2)

3. Skoulikidis NT et al. 2017 Non-perennial Mediterranean rivers in Europe: Status, pressures, and challenges for research and management. Science of the Total Environment 577, 1–18. (doi:10.1016/j.scitotenv.2016.10.147)

4. Lake PS. 2003 Ecological effects of perturbation by drought in flowing waters. Freshwater Biology 48, 1161–1172.

5. Magoulick DD, Kobza RM. 2003 The role of refugia for fishes during drought: A review and synthesis. Freshwater Biology 48, 1186–1198. (doi:10.1046/j.1365-2427.2003.01089.x)

6. Datry T, Larned ST, Tockner K. 2014 Intermittent rivers: A challenge for freshwater ecology. BioScience 64, 229–235. (doi:10.1093/biosci/bit027)

7. Dudgeon D et al. 2006 Freshwater biodiversity: Importance, threats, status and conservation challenges. Biological Reviews 81, 163–182. (doi:10.1017/S1464793105006950)

8. Su G, Logez M, Xu J, Tao S, Villéger S, Brosse S. 2021 Human impacts on global freshwater fish biodiversity. Science 371, 835–838. (doi:10.1126/science.abd3369)

9. Datry T, Bonada N, Boulton A. 2017 Intermittent Rivers and Ephemeral Streams. (doi:10.1016/b978-0-12-819166-8.00090-6)

10. Larned ST, Datry T, Arscott DB, Tockner K. 2010 Emerging concepts in temporaryCriver ecology. Freshwater Biology 55, 717–738. (doi:10.1111/j.1365-2427.2009.02322.x)

11. Leigh C, Boulton AJ, Courtwright JL, Fritz K, May CL, Walker RH, Datry T. 2016 Ecological research and management of intermittent rivers: An historical review and future directions. Freshwater Biology 61, 1181–1199. (doi:10.1111/fwb.12646)

12. Beniston M et al. 2007 Future extreme events in European climate: An exploration of regional climate model projections. Climatic Change 81, 71–95. (doi:10.1007/s10584-006-9226-z)

13. Pumo D, Caracciolo D, Viola F, Noto LV. 2016 Climate change effects on the hydrological regime of small non-perennial river basins. Science of The Total Environment 542, 76–92. (doi:10.1016/j.scitotenv.2015.10.109)

14. Mimeau L et al. 2025 Projections of streamflow intermittence under climate change in European drying river networks. Hydrol. Earth Syst. Sci. 29, 1615–1636. (doi:10.5194/hess-29-1615-2025)

15. Archdeacon TP, Reale JK. 2020 No quarter: Lack of refuge during flow intermittency results in catastrophic mortality of an imperiled minnow. Freshwater Biology 65, 2108–2123. (doi:10.1111/fwb.13607)

16. Bogan MT, Leidy RA, Neuhaus L, Hernandez CJ, Carlson SM. 2019 Biodiversity value of remnant pools in an intermittent stream during the great California drought. Aquatic Conservation 29, 976–989. (doi:10.1002/aqc.3109)

17. Vander Vorste R, Obedzinski M, Nossaman Pierce S, Carlson SM, Grantham TE. 2020 Refuges and ecological traps: Extreme drought threatens persistence of an endangered fish in intermittent streams. Global Change Biology 26, 3834–3845. (doi:10.1111/gcb.15116)

18. Kerezsy A, Gido K, Magalhães MF, Skelton PH. 2017 The Biota of Intermittent Rivers and Ephemeral Streams: Fishes. Elsevier Inc. (doi:10.1016/B978-0-12-803835-2.00010-3)

19. Davey AJH, Kelly DJ. 2007 Fish community responses to drying disturbances in an intermittent stream: A landscape perspective. Freshwater Biology 52, 1719–1733. (doi:10.1111/j.1365-2427.2007.01800.x)

20. Mas-martí E, García-Berthou E, Sabater S, Tomanova S, Muñoz I. 2010 Comparing fish assemblages and trophic ecology of permanent and intermittent reaches in a Mediterranean stream. Hydrobiologia 657, 167–180. (doi:10.1007/s10750-010-0292-x)

21. Tramer EJ. 1977 Catastrophic mortality of stream fishes trapped in shrinking pools. American Midland Naturalist 97, 469. (doi:10.2307/2425110)

22. Grossman GD, De Sostoa A. 1994 Microhabitat use by fish in the upper Rio Matarrana, Spain, 1984–1987. Ecology of Freshwater Fish 3, 141–152.

23. Kalogianni E, Vardakas L, Vourka A, Koutsikos N, Theodoropoulos C, Galia T, Skoulikidis N. 2020 Wood availability and habitat heterogeneity drive spatiotemporal habitat use by riverine cyprinids under flow intermittence. River Research & Apps 36, 819–827. (doi:10.1002/rra.3601)

24. Magoulick DD, Wilzbach MA. 1997 Microhabitat Selection by native brook trout and introduced rainbow trout in a small Pennsylvania stream. Journal of Freshwater Ecology 12, 607–614. (doi:10.1080/02705060.1997.9663575)

25. Archdeacon TP, Gonzales EJ, Yackulic CB. 2024 Fishes move to transient local refuges, not persistent landscape refuges during river drying experiment. Freshwater Biology 69, 792–808. (doi:10.1111/fwb.14246)

26. Hodges SW, Magoulick DD. 2011 Refuge habitats for fishes during seasonal drying in an intermittent stream: Movement, survival and abundance of three minnow species. Aquatic Sciences 73, 513–522. (doi:10.1007/s00027-011-0206-7)

27. Labbe TR, Fausch KD. 2000 Dynamics of intermittent stream habitat regulate persistence of a threatened fish at multiple scales. Ecological Applications 10, 1774–1791. (doi:10.1890/1051-0761(2000)010%5B1774)

28. Marshall JC et al. 2016 Go with the flow: The movement behaviour of fish from isolated waterhole refugia during connecting flow events in an intermittent dryland river. Freshwater Biology 61, 1242–1258. (doi:10.1111/fwb.12707)

29. Pires DF, Beja P, Magalhães MF. 2014 Out of pools: Movement patterns of Mediterranean stream fish in relation to ry season refugia. River Research & Apps 30, 1269–1280. (doi:10.1002/rra.2776)

30. Schiavon A, Comoglio C, Candiotto A, Spairani M, Hölker F, Tarena F, Watz J, Nyqvist D. 2024 Navigating the drought: Upstream migration of a small-sized Cypriniformes (*Telestes muticellus*) in response to drying in a partially intermittent mountain stream. Knowl. Manag. Aquat. Ecosyst., 6. (doi:10.1051/kmae/2024003)

31. Bonada N et al. 2020 Conservation and management of isolated pools in temporary rivers. Water 12, 2870. (doi:10.3390/w12102870)

32. Hopper GW, Gido KB, Pennock CA, Hedden SC, Frenette BD, Barts N, Hedden CK, Bruckerhoff LA. 2020 Nowhere to swim: interspecific responses of prairie stream fishes in isolated pools during severe drought. Aquatic Sciences 82, 1–15. (doi:10.1007/s00027-020-0716-2)

33. Magoulick DD. 2000 Spatial and temporal variation in fish assemblages of drying stream pools: The role of abiotic and biotic factors. Aquatic Ecology 34, 29–41. (doi:10.1023/A:1009914619061)

34. Pires DF, Pires AM, Collares-Pereira MJ, Magalhães MF. 2010 Variation in fish assemblages across dry-season pools in a Mediterranean stream: Effects of pool morphology, physicochemical factors and spatial context. Ecology of Freshwater Fish 19, 74–86. (doi:10.1111/j.1600-0633.2009.00391.x)

35. Schiavon A, Comoglio C, Candiotto A, Spairani M, Hölker F, Watz J, Nyqvist D. 2025 Individual movement behaviour and habitat use of a small-sized cypriniform (*Telestes muticellus*) in a mountain stream. Environ Biol Fish 108, 241–258. (doi:10.1007/s10641-024-01661-9)

36. Ayllón D, Almodóvar A, Nicola GG, Elvira B. 2010 Ontogenetic and spatial variations in brown trout habitat selection. Ecology of Freshwater Fish 19, 420–432. (doi:10.1111/j.1600-0633.2010.00426.x)

37. Harvey BC, Stewart AJ. 1991 Fish size and habitat depth relationships in headwater streams. Oecologia 87, 336–342. (doi:10.1007/BF00634588)

38. Vardakas L, Kalogianni E, Papadaki C, Vavalidis T, Mentzafou A, Koutsoubas D, Skoulikidis Th. N. 2017 Defining critical habitat conditions for the conservation of three endemic and endangered cyprinids in a Mediterranean intermittent river before the onset of drought. Aquatic Conservation 27, 1270–1280. (doi:10.1002/aqc.2735)

39. Stoffers T, Buijse AD, Poos JJ, Verreth JAJ, Nagelkerke LAJ. 2025 Ontogenetic shifts by juvenile fishes highlight the need for habitat heterogeneity and connectivity in river restoration. Limnology & Oceanography 70, 732–748. (doi:10.1002/lno.12797)

40. Gasith A, Resh VH. 1999 Streams in Mediterranean climate regions: Abiotic Influences and Biotic Responses to Predictable Seasonal Events. Annual Review of Ecol Syst 30, 51–81.

41. Hermoso V, Clavero M. 2011 Threatening processes and conservation management of endemic freshwater fish in the Mediterranean basin: A review. Marine and Freshwater Research 62, 244–254. (doi:10.1071/MF09300)

42. Pou-Rovira Q, Ferrer D, Cruset E, Sànchez S, Arquimbau R. 2013 Diagnosi de les poblacions de peixos i dels seus hàbitats a l’espai natural de l’Alta Garrotxa. Annals de la delegació de la Garrotxa de la Inst. Cat. Hist. Nat. 6, 77–86.

43. Doadrio I, Perea S, Garzón-Heydt P, González JL. 2011 Ictiofauna Continental Española.

44. Aparicio E. 2016 Peixos continentals de Catalunya:Ecologia, conservació i guia d’identificació. Lynx Edicions.

45. Merciai R, Molons-sierra C, Sabater S, García-Berthou E. 2017 Water abstraction affects abundance, size-structure and growth of two threatened cyprinid fishes. PloS one

46. Merciai R, Bailey LL, Bestgen KR, Fausch KD, Zamora L, Sabater S, García-Berthou E. 2018 Water diversion reduces abundance and survival of two Mediterranean cyprinids. Ecology of Freshwater Fish 27, 481–491. (doi:10.1111/eff.12363)

47. Ford M. 2024 Barbus meridionalis. In The IUCN Red List of Threatened Species 2024, p. e.T2567A137226182.

48. Wilson ADM, Krause S, Ramnarine IW, Borner KK. 2015 Social networks in changing environments. Behavioral Ecology and Sociobiology 69, 1617–1629. (doi:10.1007/s00265-015-1973-2)

49. Akima H, Gebhardt A, Petzold T, Maechler M. 2016 Package ‘akima’. Version 0.6 2.

50. Jolles J. 2026 ATracker: A flexible Python toolkit for animal tracking. (doi:10.5281/zenodo.18889019)

51. Schiavon A, Comoglio C, Candiotto A, Spairani M, Hölker F, Watz J, Nyqvist D. 2024 River runs dry: Movement patterns of *Telestes muticellus (Cypriniformes: Leuciscidae*) in an intermittent river stretch. In Advances in Hydraulic Research (eds MB Kalinowska, MM Mrokowska, PM Rowiński), pp. 341–351. Cham: Springer Nature Switzerland. (doi:10.1007/978-3-031-56093-4_27)

52. Aparicio E, De Sostoa A. 1999 Pattern of movements of adult *Barbus haasi* in a small Mediterranean stream. Journal of Fish Biology 55, 1086–1095. (doi:10.1111/j.1095-8649.1999.tb00743.x)

53. Santos JM, Rivaes R, Boavida I, Branco P. 2018 Structural microhabitat use by endemic cyprinids in a MediterraneanCtype river: Implications for restoration practices. Aquatic Conservation 28, 26–36. (doi:10.1002/aqc.2839)

54. Cunjak RA, Green JM. 1983 Habitat utilization by brook char (*Salvelinus fontinalis*) and rainbow trout (*Salmo gairdneri*) in Newfoundland streams. Can. J. Zool. 61, 1214–1219. (doi:10.1139/z83-165)

55. Santos JM, Ferreira MT. 2008 Microhabitat use by endangered Iberian cyprinids nase *Iberochondrostoma almacai* and chub *Squalius aradensis*. Aquat. Sci. 70, 272–281. (doi:10.1007/s00027-008-8037-x)

56. Espírito-Santo HMV, Rodríguez MA, Zuanon J. 2017 Strategies to avoid the trap: Stream fish use fine-scale hydrological cues to move between the stream channel and temporary pools. Hydrobiologia 792, 183–194. (doi:10.1007/s10750-016-3054-6)

57. Magalhães MF, Beja P, Canas C, Collares-Pereira MJ. 2002 Functional heterogeneity of dry-season fish refugia across a Mediterranean catchment: The role of habitat and predation. Functional Biology 47, 1919–1934.

58. Schiavon A, Comoglio C, Candiotto A, Spairani M, Hölker F, Watz J, Nyqvist D. 2025 Movement pattern and habitat use of the endangered brook Barbel (*Barbus caninus*) in a Mediterranean stream. Ecology of Freshwater Fish 34, e70017. (doi:10.1111/eff.70017)

59. Wong BBM, Candolin U. 2015 Behavioral responses to changing environments. Behavioral Ecology 26, 665–673. (doi:10.1093/beheco/aru183)

60. Hermoso V, Ward DP, Kennard MJ. 2013 Prioritizing refugia for freshwater biodiversity conservation in highly seasonal ecosystems. Diversity and Distributions 19, 1031–1042. (doi:10.1111/ddi.12082)

