## Supplementary figure for "Fine-scale movements and habitat use of fish in intermittent rivers: Behavioural insights from drying refuge pools"

*For*

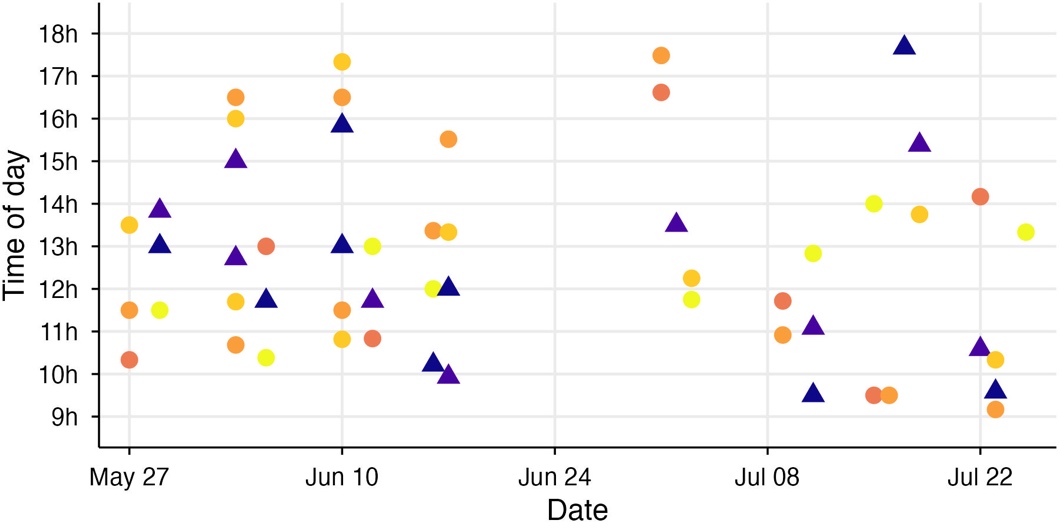

***Figure S1.*** *Temporal distribution of observation sessions across the monitoring period. Each point indicates the start time of an observation session on a given date, illustrating pseudo-random sampling across days and times of day. Drying pools are shown in orange-yellow tones with circular symbols, and non-drying pools in dark blue with triangular symbols.*

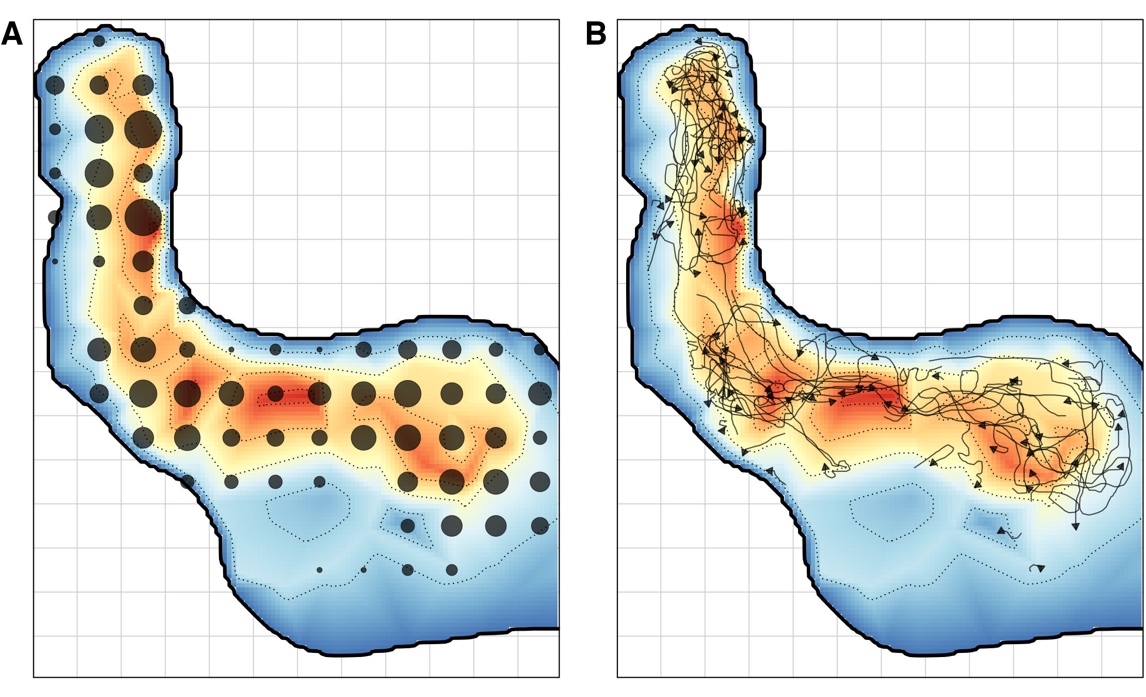

***Figure S2.*** *Example spatial distribution of behavioural observations within a representative pool (DP3). (A) Grid-based observations showing fish detections across 1 m × 1 m grid cells. Circle size reflects the frequency of detections across surveys. (B) Focal-follow trajectories of individual fish overlaid on the same pool. Background colours indicate relative water depth derived from the interpolated depth surface; contour lines represent 10 cm depth intervals.*

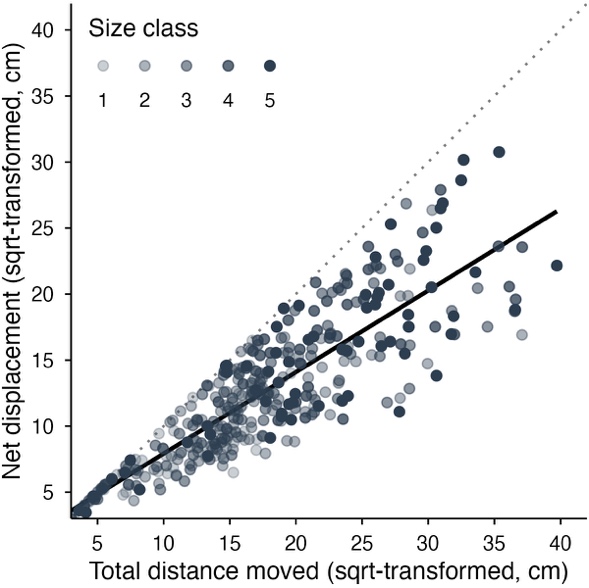
***Figure S3.*** *Relationship between total and maximum displacement across focal follows (square-root scale). The dashed diagonal indicates a perfectly straight movement path, whereas the solid black line shows the fitted relationship used to quantify path straightness. Deviations from this line represent residuals, with values above the line indicating straighter-than-expected trajectories and values below indicating more tortuous movement. For analysis, residuals were scaled by maximum displacement to obtain a relative path-straightness metric.*

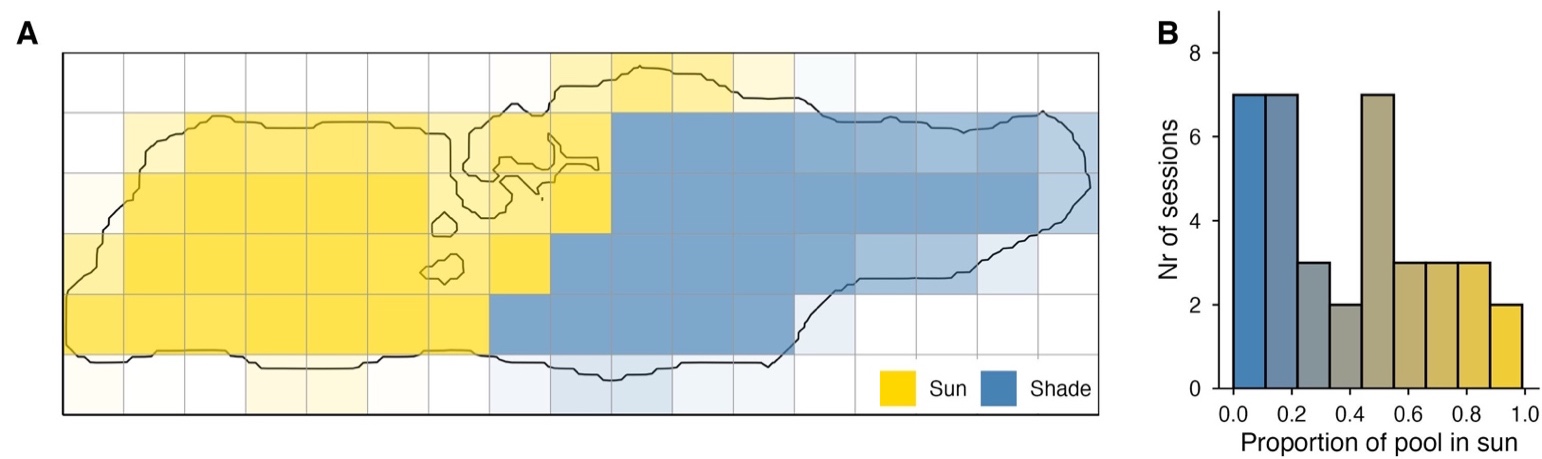

***Figure S4.*** *(A) Example sun-exposure map from one grid-observation session (NP2). Each square represents a 1 m² grid cell, classified as receiving direct sunlight (yellow) or being in the shade (blue). Black outline indicates the pool waterline at the time of observation. (B) Distribution of the proportion of wet grid cells receiving direct sunlight across all monitoring sessions; the 13 sessions without any sun are not shown. Bars are coloured blue to yellow to indicate increasing sun exposure*

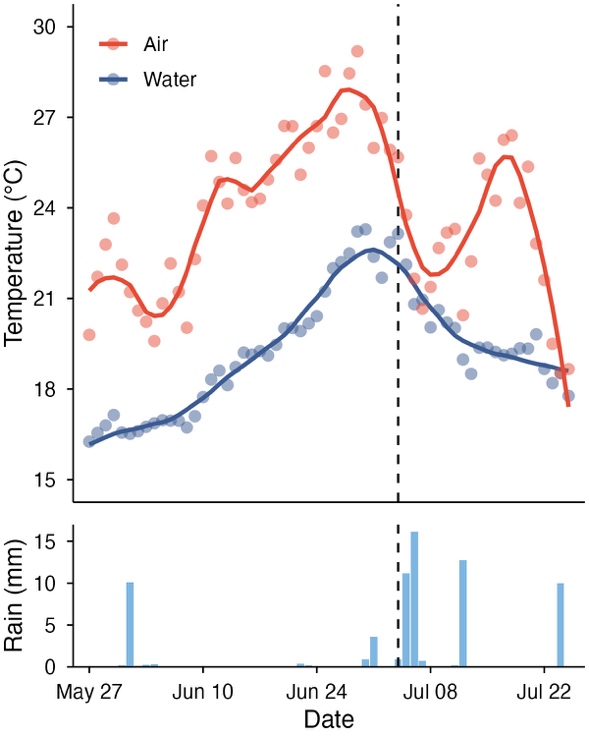

***Figure S5.*** *Temporal variation in temperature and rainfall across the study period. Daily air temperature (red) and water temperature in a representative pool (blue) are shown in the upper panel, with rainfall totals in the lower panel. Points represent observed daily values and solid lines indicate LOESS-smoothed trends. The dashed vertical line marks the transition from the main drying phase to the subsequent refilling period.*

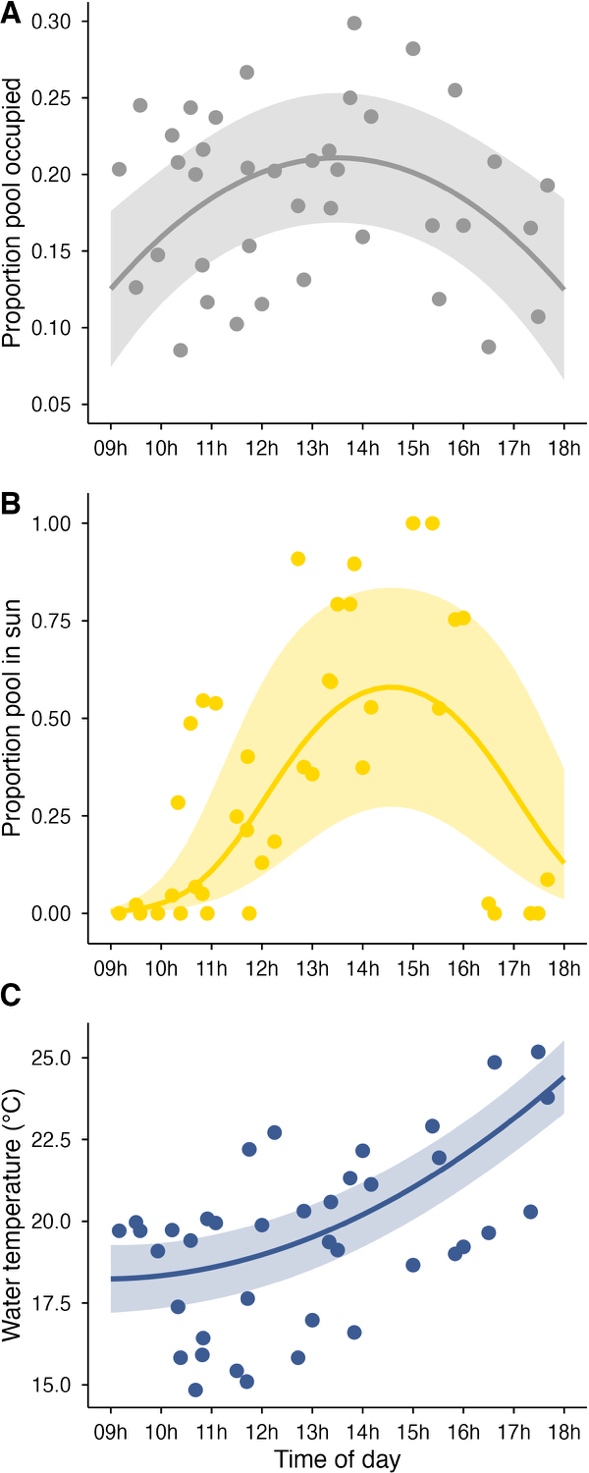

***Figure S6.*** *(A) Predicted diurnal pattern in pool occupancy from a mixed-effects model including a second-order polynomial of time of day. (B) Diurnal variation in the proportion of wet grid cells receiving direct sunlight, shown as model-based predictions from a binomial mixed-effects model with a quadratic time term. (C) Diurnal variation in pool water temperature, shown as predictions from a linear mixed-effects model with a quadratic effect of time of day. Shown are means ± 95% confidence intervals.*

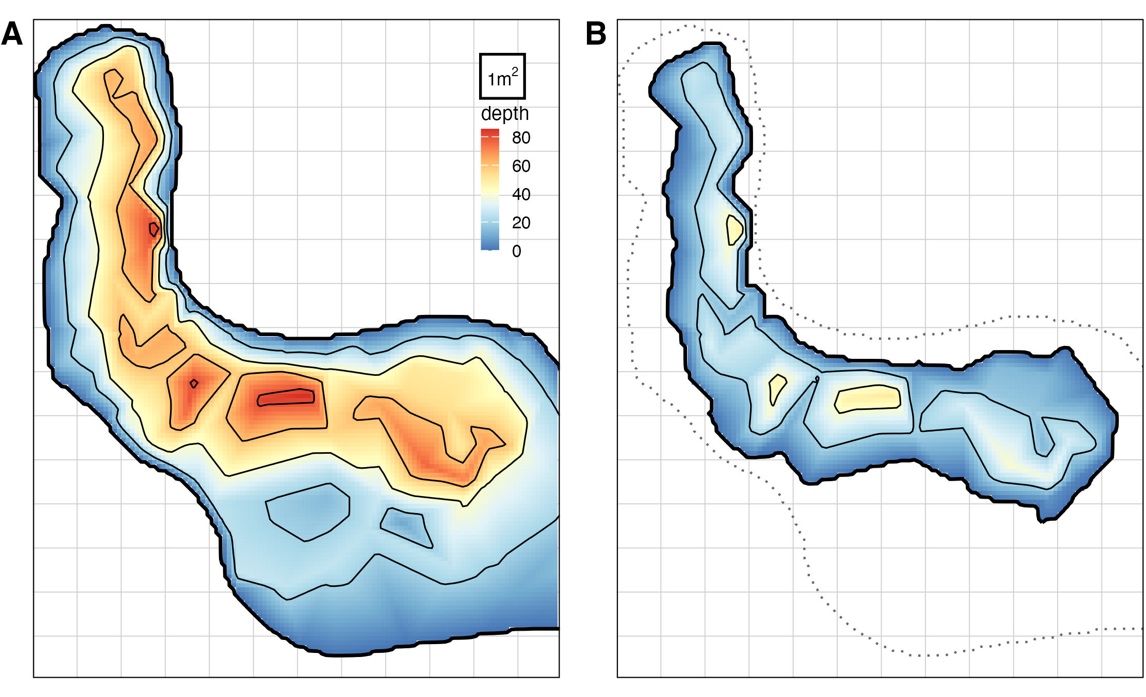

***Figure S7.*** *(A) Pool bathymetry at the start of the monitoring period, prior to substantial drying, when water level and surface area were maximal. (B) Bathymetry of the same pool at the peak of the drying phase, when water level reached its seasonal minimum. The grid matches that used for the grid observations, as depicted by the bordered grid cell. Colours indicate water depth (cm) derived from an interpolated depth grid at 10 × 10 cm resolution, with contour lines spaced at 10 cm depth intervals. In panel B, the dotted outline indicates the maximum pool extent earlier in the season, illustrating contraction of pool area during drying.*

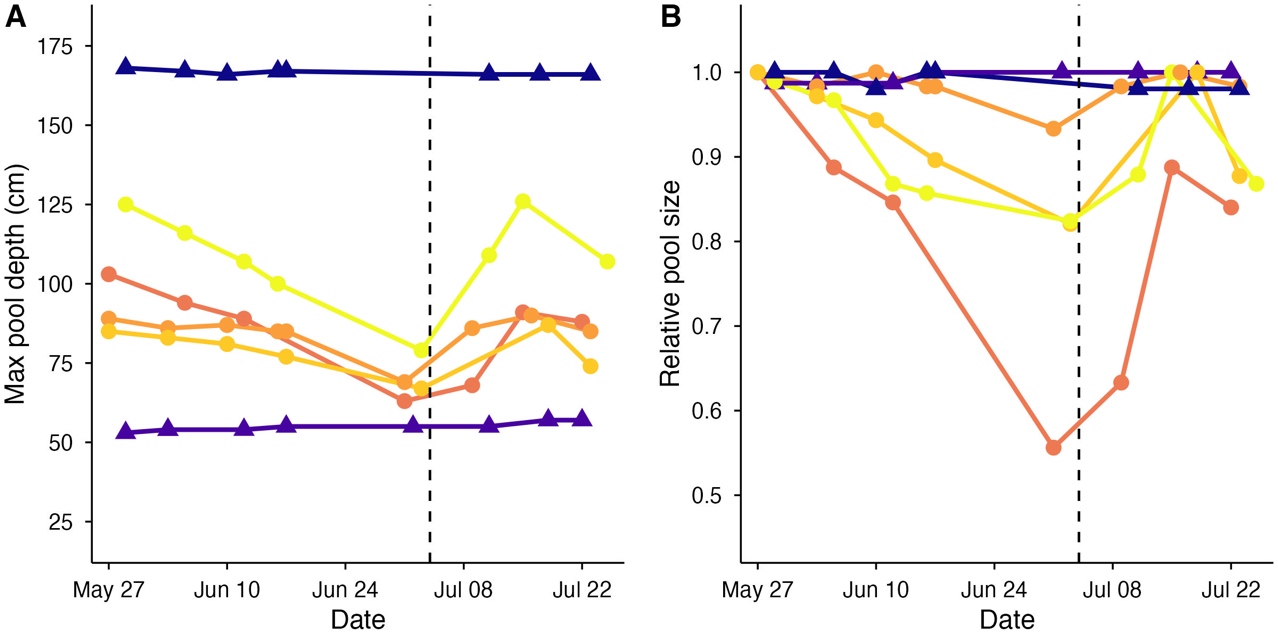

***Figure S8.*** *(A) Maximum pool depth and (B) pool surface area, expressed relative to the maximum observed size during the study period, for each monitored pool. Individual pools are shown as coloured lines; drying pools are indicated in orange–yellow with circles, and non-drying pools in dark blue with triangles. The dashed vertical line marks the transition between the main drying phase and the subsequent refilling period used in the analyses.*
